# A sustainable and bee-pollinated coffee to start your day

**DOI:** 10.1101/2024.09.30.615918

**Authors:** Denise Araujo Alves, Jenifer Dias Ramos, Charles Fernando dos Santos, Diego Moure-Oliveira, Joyce Mayra Volpini Almeida-Dias, Gustavo Souza Santos, Thamires Sá de Oliveira Kaminski, Ana Paola Cione, Fernando Celso Longhim Quenzer, Alistair John Campbell, Andresa Aparecida Berretta, Guilherme Jorge Gomes e Sousa, Andrigo Monroe Pereira, Helen Thompson, Natalia Bottan de Toledo Vasconcelos, Rafael Cruz, Ana Carolina Martins de Queiroz, José Maurício Simões Bento, Cristiano Menezes

## Abstract

The integration of managed pollinators into agricultural systems represents a practical strategy for boosting crop production, optimising sustainable farming practices, and improving rural livelihoods. However, the use of managed bees to supplement the contribution of wild pollinators still faces challenges, such as over-reliance on pesticides in intensively managed agroecosystems. Here, we placed managed bee colonies on conventional farms to assess their impact on arabica coffee yield, quality and market value, while also to evaluate colony health in coffee fields treated with field rates of thiamethoxam-based products. We show that supplementing farms with managed bees increased farmer income by enhancing coffee yield and beverage quality, which is rewarded by specialty coffee markets. Moreover, our field-based study reveals that, even when exposed to field-realistic levels of the neonicotinoid thiamethoxam, managed bee colonies exhibited no significant adverse effects on their health within coffee fields. Our results underline that pollination is a key input on intensively managed coffee farms, and that agricultural production and environmental conservation interact synergically to maximize profitability, which is fundamental to encourage farmers to be good stewards of their croplands by adopting nature-positive practices.

## Main

Providing an adequate supply of agricultural crops for a world population projected to reach nearly 10 billion people by 2050, with minimal environmental and public health impacts, poses a significant transformative challenge^1,2^. Although substantial efforts to increase crop yields and economic outcomes could potentially reduce the environmental footprint of agriculture, more ambitious sustainability-oriented practices are needed to achieve resilient and productive farming systems^3^. The transition towards more sustainable practices has considerable positive outcomes, providing food and nutrition security, improving profitability, reducing social inequality, and safeguarding ecosystem goods and services on which agriculture relies^3,4^, such as pollination. Animal mediated-pollination services are important not only for yield increase, but also enhance aesthetic aspects, nutritional composition, shelf lifetime, and market value of several crops, ultimately contributing to increased profitability for farmers^5–9^.

Over past decades, while pollinator-dependent crops have expanded substantially^10^, the richness and abundance of wild and managed pollinators have declined due to multiple interconnected human-induced pressures, including pesticide misuse, land-use changes, and climate change^7,11^. These drivers have imposed constraints on the quantity and stability of agricultural production, particularly in tropical regions^12^. Although the majority of pollinator-dependent crops are grown in the tropics, more developed countries rely on pollinated crop imports to sustain their consumption patterns^13^. Consequently, these countries may suffer the impacts of pollinator decline in their trade partners^13^. Among the most valuable tropical pollinator-dependent cash crops, coffee is an integral part of the global supply chain, contributing billions of dollars annually^14^. Coffee stands as the second most consumed beverage globally, appreciated for its pleasurable qualities and health benefits^15^, and its commodity chain directly connects producers, from small to large-sized farms, with coffee drinkers in urban centres^16^. Moreover, growing consumer interests in superior beverage quality and environmentally sound products have pushed farmers to promote higher sustainability standards in coffee systems^15,16^.

Considering that bee pollination significantly enhances fruit set and weight^17,18^ of the higher-quality arabica coffee (*Coffea arabica*), a self-fertile crop, pollinator loss due to climate change and agricultural land use has raised concerns about its negative consequences for coffee productivity^12^, which supports millions of rural livelihoods across the tropics^16^. Thus, in a changing world, the integration of managed pollinators into agricultural systems is essential to ensure stable pollination services and optimise sustainable farming practices^19,20^. However, the deployment of managed bees to supplement the contribution of wild pollinators still faces challenges^20^. One major hurdle is the over-reliance on chemical pesticides to reduce pest damage and protect crop quality in intensively managed agroecosystems. Depending on the mode of action and scale of exposure to these pesticides, or their environmental metabolites, foraging on treated crops may result in lethal or trigger subtle effects on behavioural and physiological traits of social bees^21^, including foraging performance, learning and memory, colony development, and disease resistance, which may ultimately compromise their health^7,21,22^. Also, some chemical pesticides are commonly found in food stores of bee nests^23^, thereby extending the exposure of social bees. Although neonicotinoids, such as thiamethoxam, are efficient in controlling economically important crop pest populations, there has been global concern about their harmful effects on pollinators due to their widespread use, persistence and high systemicity, meaning that they are absorbed by the treated plant and translocated to all plant organs, reaching nectar and pollen in flowers^24^. To address risks to bees, many countries have national pesticide regulation, which requires studies from laboratory to field scales, using primarily the honeybee *Apis mellifera* as a surrogate for thousands of bee species^25^. Given the diversity in life history and physiology, growing evidence shows that there is a substantial interspecific variation in pesticide susceptibilities, highlighting the need to (re)assess lethal doses and/or exposure routes of the pesticide products that could have relevant impacts on non-*Apis* bees^25,26^, such as the tropical social stingless bees^26,27^. For instance, stingless bees may be threatened by pesticides that are harmless to *A. mellifera*, as most species rely on symbiont microorganisms and their biomolecules to prevent stored food from spoiling and to improve its nutritional/hormonal quality^28^, or even *Melipona* species that depend on mud to build their nests^29^. Furthermore, most ecotoxicological studies are focused on lethal effects at the individual level, while sub-lethal effects on a range of behavioural traits that are crucial for colony growth and functioning of social bees, especially under field conditions, have received less attention^21^.

In the context that pollinators play a crucial role for crop production and human well-being^5–7^ and that coffee is an outstanding potential crop to achieve a “win-win” scenario in transition pathways towards environmentally and economically sustainable agriculture^30^, we addressed three hypotheses: (H1) the supplemental pollination provided by managed social bees significantly improves both coffee yields and (H2) quality, promoting monetary benefits to farmers, and (H3) the use of neonicotinoids at field rates, based on the pest control efficiency, results in acceptable risk to colony bee health within conventional coffee farming systems. Despite the evidence on the benefits of pollination to arabica coffee production and quality^17,18,31,32^, to our knowledge, ours is the first study to consistently show that supplementation of farms with managed bees results in higher farmers’ economic outcomes by enhancing coffee productivity and beverage quality, which is rewarded by specialty coffee markets. In addition, our field-based study provides significant new evidence on the effects of pesticide exposure on tropical bee colonies. We demonstrate that under realistic field settings, there is no significant effect on the bee colony health when managed in coffee fields treated with field rates of thiamethoxam-based products.

We addressed these issues through manipulative experiments at realistic coffee field scales. To achieve this, we introduced healthy bee colonies, the exotic Africanised honeybee *Apis mellifera* (in 2021/2022) and the native stingless bee *Scaptotrigona depilis* (in 2022/2023), on intensively managed coffee-producing farms in Southeastern Brazil. For each farm, we assessed coffee yield per hectare (*A. mellifera*) and per branch (*S. depilis*) in coffee sites near to bee colonies (i.e., within 50 m, hereafter, ‘with managed bees’) and in more distant control sites (i.e., mean distance of 250 m from bee colonies, hereafter, ‘without managed bees’). By combining chemical, physical, and sensory analyses and following the same experimental design, we examined the effects of managed honeybees on coffee quality and market value. Finally, we evaluated the effect of thiamethoxam and its metabolite clothianidin residues on stingless bees by assessing their colony performance on both conventionally managed and pesticide-free control coffee-producing farms (hereafter, conventional group and organic group, respectively; Methods).

## Results and Discussion

Our manipulative field study shows that pollination provided by managed social bees significantly increased arabica coffee yield both at the field and shrub levels (H1, Fig. 1). At the field level, placing honeybee colonies on conventional farms increased the mean number of coffee bags per hectare by 16.5% (mean ± SE; with *vs*. without managed bees: 37.90 ± 4.813 bags ha^-1^ *vs*. 32.51 ± 4.262 bags ha^-1^; Fig. 1a and Extended Data Table 1), confirming our previous finding that coffee yields per shrub increased by an average of 16% near managed honeybees^33^. At the shrub level, the deployment of managed stingless bees resulted in a significant increase of 67% in weight of fruit set per branch (with *vs*. without managed bees: 0.50 ± 0.002 kg branch^-1^ *vs*. 0.30 ± 0.008 kg branch^-1^; Fig. 1b and Extended Data Table 1). It should be noted that differences in experimental designs, harvesting methods, blooming seasons, and edaphoclimatic conditions between the conventionally managed coffee farms complicate comparisons of pollination performance among honeybees and stingless bees. Indeed, despite it was not our aim to contrast the effect of the managed bee species on coffee yield, as we did not supplement farms with stingless bee colonies along with honeybees, our data provide useful baseline figures for designing adequate experimental setups to test this hypothesis.

**Fig. 1.**
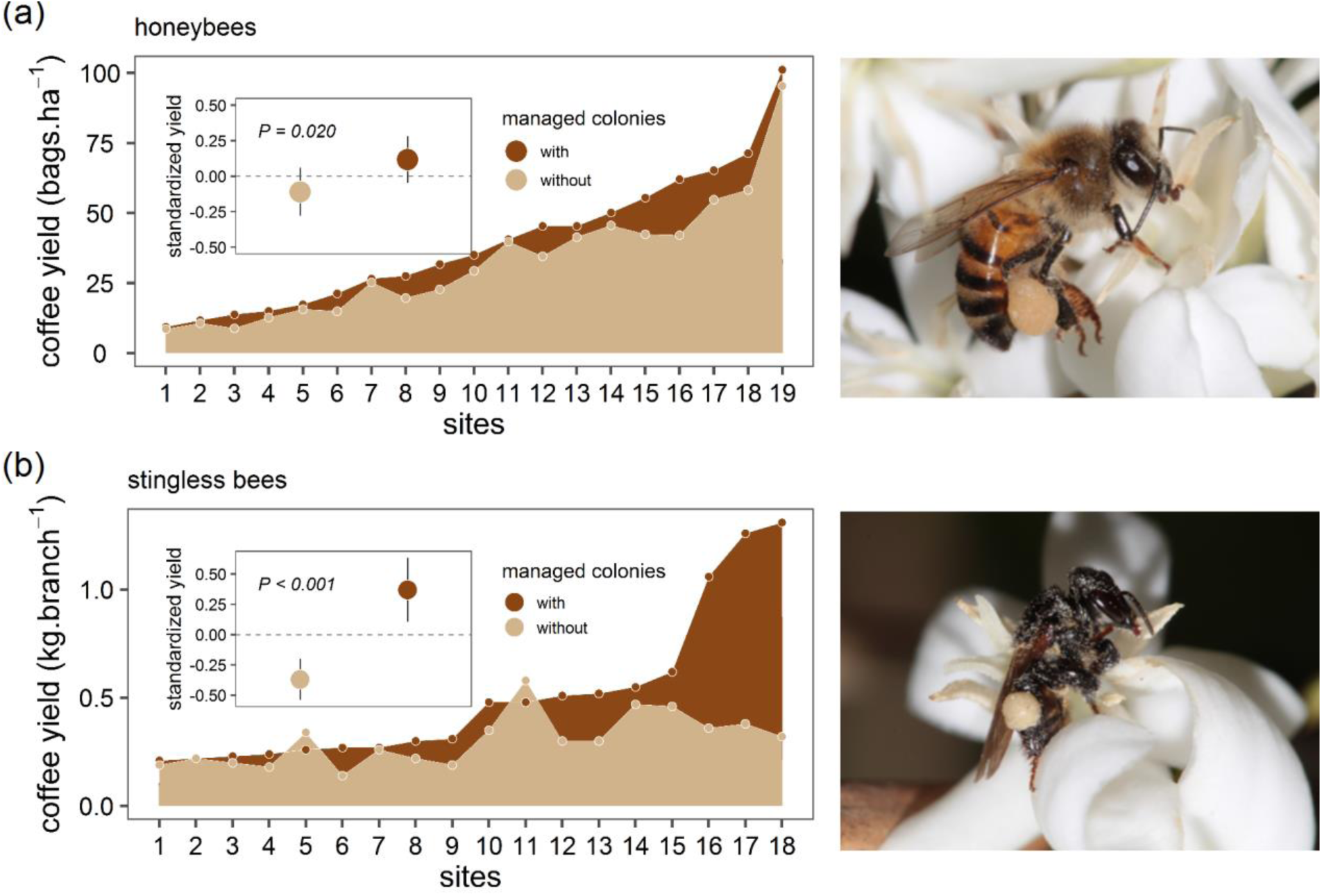
| Benefits of managed bees to coffee yield. Increased coffee yield (bags ha^-1^ and kg branch^-1^) with the deployment of managed (**a**) honeybee and (**b**) stingless bee colonies on coffee sites in dark brown. Coffee yields in fields without managed bee colonies are in light brown. Inner figures correspond to the effect of pollination treatment (with and without managed bees) on log-transformed coffee yield (data are standardized with mean 0 ± SE to allow comparison of effect sizes).

Although it is well known that honeybees and stingless bees are the most important coffee flower visitors at the field and shrub levels^34^, managed pollination practices to increase yields are largely focussed on augmenting the exotic honeybee abundance by placing hives on farms. Nevertheless, our results support that the native *Scaptotrigona* stingless bee, with typical shorter foraging range (∼0.9 *vs*. ∼1.3 km)^35,36^ and narrower diet breath than the supergeneralist honeybees^37^, is an highly effective managed pollinator for coffee production in the Neotropical Region^33^. The high pollination efficiency of *S. depilis* is likely explained by its morphological and behavioural traits, including small body size (∼5.5 mm), short trip duration (∼4.2 min), constant foraging activity from morning to early afternoon, mass-recruitment foraging strategy using scent trails to quickly mobilise nestmates to food sources, and large colony size (∼10,000 workers)^29^. In addition, although *S. depilis* foragers are similar to honeybees in moistening the collected coffee pollen with nectar and compacting it into their metatibial pollen baskets, apparently, they carry more free pollen grains incidentally attached to their bodies that are more likely to be available for pollination (Fig. 1). Even though stingless bees are well suited as managed tropical pollinators, as they pose less of a hazard to farm workers due to the lack of a functional sting and have relatively well-developed colony management protocols, commercial colony production is still incipient^38^ compared with large-scale beekeeping operations, which provides hundreds of honeybee colonies to support crop pollination. Even so, the shortage in pollinator availability is a common problem in many agroecosystems, rising pollination deficits and reducing crop yield^39^. At the same time, there is a growing concern over reliance on *A. mellifera* as a single key species for agricultural pollination, mainly in regions outside its native range, which has increased interest in native bees as alternative managed pollinators to optimise crop pollination^20^.

Beyond increased coffee yield by 16.5%, supplemental pollination provided by managed honeybees significantly improved coffee quality, promoting monetary benefits to farmers (H2, Fig. 2). Our data show a significant increase in the coffee quality parameters in areas with managed honeybees (PA; M^2^ = 0.19, R = 0.89, *P* = 0.043). Overall, some farms supplemented with honeybee colonies showed substantial displacements in the Procrustes space (i.e., with significant impact on coffee quality attributes), while others maintained a reasonable level of similarity. This is evident in the Procrustes residuals’ position, length, and direction, which seem to influence coffee flavour and score (Fig. 2a). Our findings highlight the complex relationship between the supplemental pollination mediated by managed honeybees and coffee quality parameters. Moreover, our data show that the deployment of managed honeybees significantly improved coffee score by 2.43 points (GLM, χ^2^ = 5.14, *P* = 0.02; Fig. 2b). Two out of six farms experienced beverage classification enhancement when managed honeybees were deployed – from below to very good specialty coffee and from very good to excellent specialty coffee (Fig. 2a) –, enabling farmers to make profits from the higher prices of the specialty coffee market. Indeed, since coffee market value was based on coffee score and bean size ≥16, rated as exportable beans, pollination provided by managed honeybees significantly improved economic benefit by 13.15%, which means a surplus of US$ 25.36 per coffee bag for farmers (GLM, χ^2^ = 4.19, *P* = 0.04; Fig. 2c). Although there was not a significant effect of supplemental pollination provided by managed honeybees on several coffee quality parameters, such as the amount of volatile compounds^31^, it still slightly improved bean size by 2.14% and reduced the percentage of peaberries (i.e., misshapen berries with one seed) by 9.71% and caffeine content by 2.78% (Fig. 2d-f). Since a high level of caffeine is associated with low quality brews^40^, even small reductions of this bioactive compound in roasted beans might modify the bitter and astringent flavour of the brewed coffee.

**Fig. 2.**
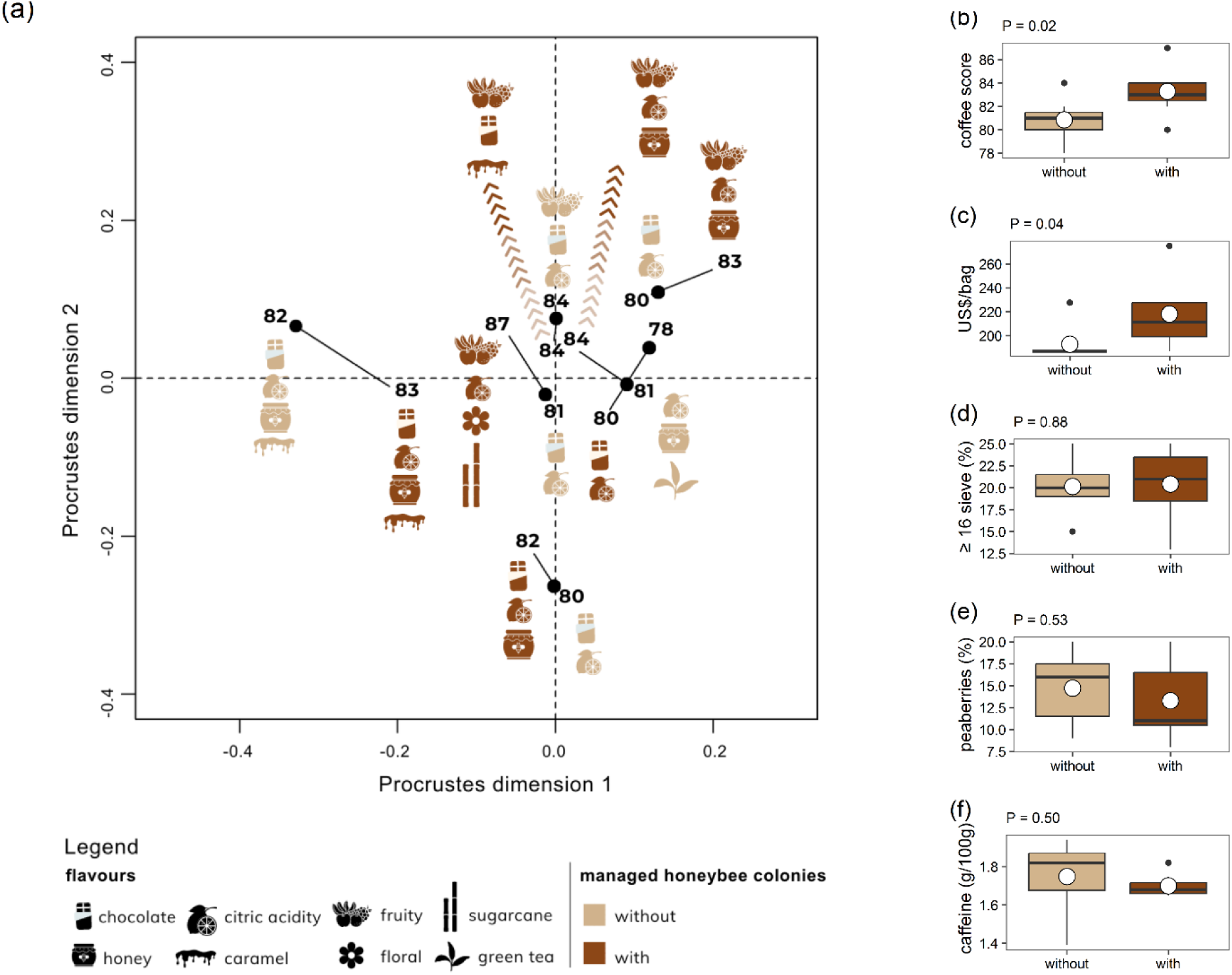
| Contribution of managed honeybees to coffee quality. (**a**) Significant mismatch (Procrustes analysis, M^2^ = 0.19, R = 0.89, *P* = 0.04) between the coffee fields with and without managed honeybee colonies, based on 44 coffee parameters (physical, chemical, sensorial, and economic). These discrete shifts are significant enough for coffee beverage quality (scores and flavour) improvement on fields with managed bees (initial position of each field without managed bees is represented in light brown, while the corresponding displacements towards to field with managed bees are in dark brown). The trajectories and the residual length (straight lines) indicate the degree of discrepancies between fields with and without managed bees after achieving an optimal fit. (**b-f**) Key coffee parameters in line with market interests: (**b**) beverage classification on a 100-point scale, (**c**) market value of 60 kg bags for ≥16-sieved coffee beans, percentage of (**d**) ≥16-sieved coffee beans, (**e**) percentage of peaberries, and (**f**) caffeine content. The boxplots in (**b**)**-**(**f**) show the mean (circle), the median (horizontal line), the 25-75th percentile range (box), and the data distribution (whisker).

As expected by previous studies, pollinators contribute to arabica coffee quality by improving yield^17,18,32^, physical parameters (e.g., reduced percentage of peaberries^17^ and increased bean size^18,41^), as well as cup quality^41^ and economic value^17,18,32^. Nevertheless, our study is unique in that we manipulated pollinator abundance in coffee fields, rather than assessed the effect of pollination services provided by wild insects on coffee quality^18,31,32,41^. Furthermore, we carried out a comprehensive evaluation of a large number of parameters influencing coffee quality that might be affected by supplementation of managed honeybees and, ultimately, its market value. Even though we did not detect significant effects of managed bees for many of the coffee parameters, our results are surprising given that our experimental design contrasted the vast majority of previous studies, which used pollinator exclusion experiments and showed increased coffee production in the presence of naturally occurring wild pollinators^19,33,34,46^. Moreover, we demonstrated that by artificially augmenting pollinator abundance through bee nest management coffee farmers raise their profitability. Overall, our results provide robust evidence that the use of managed bees, in an integrated approach to pollination management^20^, boosts not just coffee productivity but also beverage quality, representing real economic benefits for coffee farmers, especially those involved in specialty coffee markets^16,42^.

Here, we show that supplemental pollination provided by managed bees is undoubtedly a key input into intensively managed coffee-producing farms. As honeybees play a significant role for improvement of arabica coffee yield and quality, at field scale, integrate them with multiple managed native bee species would maximize provision of pollination services and economic output^20,39^ for coffee farmers. Such approach to promote managed multi-species on coffee production systems in the short-term should be integrated with long-term habitat enhancement strategies to diversify wild pollinator communities, through conservation or restoration of natural habitats^19,20^ within coffee farms^43,44^. Even that associated costs of restoration programs act as the main financial barrier for their adoption, in the long-term, restoring degraded lands within coffee farm systems provides multiple functional benefits to farmers and society^43,44^, including low costs associated with supplementation of farms with managed colonies. In the Atlantic Forest, the same biome of 7 out of 23 conventional farms where we conducted our study, over 20 years, restoration costs within coffee farms can be offset by both boosting coffee yields through pollination and earnings from carbon sequestration, when farms had at least 10% forest cover at the start of the restoration process^47^. Therefore, the effective implementation of restoration efforts in coffee lands promotes conservation of biodiversity and ecosystem services, climate mitigation, increased coffee productivity and economic gains^47^.

To optimise pollination services, one of the combined tactics for a successful integrated pollinator and pest management approach is enhancing the farm environment through pesticide stewardship to mitigate pesticide risks for non-target beneficial insects^20^. Honeybees are used as model species for environmental monitoring and risk assessment, but non-*Apis* bee species, such as stingless bees, are relatively neglected^21,26^. Given that the diversity of pollinators has consistently been shown to be more relevant than the abundance of a single bee species for crop pollination^7,19^, understanding the underlying factors driving exposure in different bee species is essential for developing strategies to mitigate pesticide risks^27^.

Our field-scale study shows that the use of field rates of thiamethoxam-based products, applied by soil drenching on arabica coffee farms, had no negative relevant effects on managed stingless bee colony health (H3, Fig. 3). Managed stingless bee colonies kept in conventional farms showed the same pattern of brood production when compared to those colonies kept in organic farms (GLMM, quasi-Poisson distribution, χ^2^ = 2.61, *df* = 1, *P* = 0.10; Fig. 3a) or their interaction (GLMM, χ^2^ = 4.69, *df* = 4, *P* = 0.32). However, brood production differed among periods within each farming system (GLMM, χ^2^ = 81.1, *df* = 4, *P* < 0.001; Supplementary Table 3). The same pattern was found for brood mortality. While the type of farming system did not affect the brood mortality (GLMM, negative binomial distribution, χ^2^ = 0.20, *df* = 1, *P* = 0.65; Fig. 3b), or the interaction between them (GLMM, χ^2^ = 6.97, *df* = 4, *P* = 0.13), the brood mortality rate differed among periods within the farming system (GLMM, χ^2^ = 68.6, *df* = 4, *P* < 0.001; Supplementary Table 4). On the other hand, we detected significant differences in foraging activity over time (GLMM, quasi-Poisson distribution, χ^2^ = 70.3, *df* = 4, *P* < 0.001; Fig. 3c), between farming systems (GLMM, χ^2^ = 7.2, *df* = 1, *P* = 0.006) and the interaction between them (GLMM, χ^2^ = 29.3, *df* = 4, *P* < 0.001; Supplementary Table 5).

**Fig. 3.**
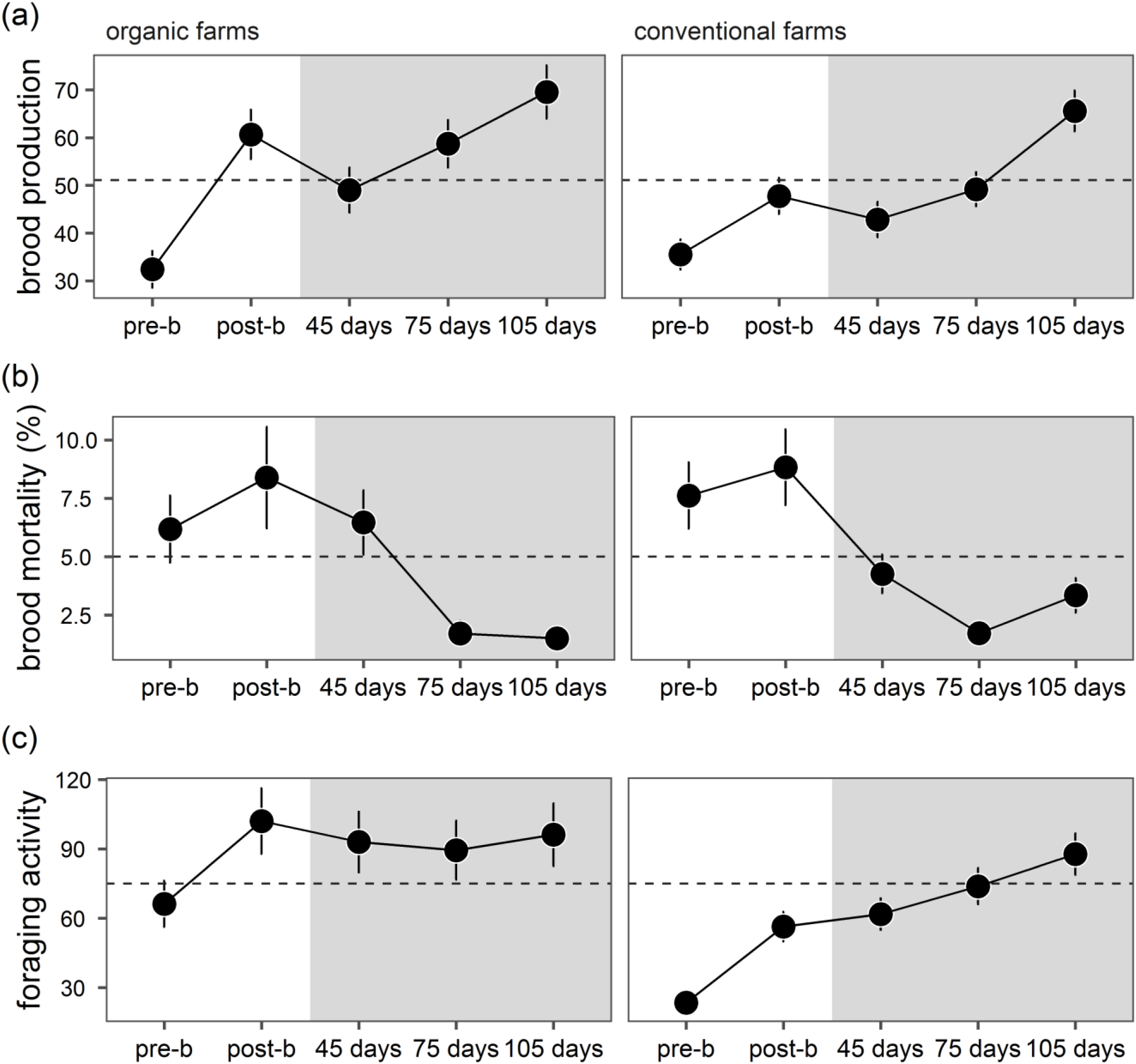
| Effects of thiamethoxam-based products on stingless bee colony health in coffee farms. Brood production (**a**), brood mortality (**b**) and foraging activity (**c**) were assessed on both conventionally managed (right) and chemical pesticide-free control (left) farms in five periods: pre- and post-blooming and the subsequent 45, 75, and 105 days. Evaluations were carried out at coffee farms (in white) and at a forest patch located over 90 km south of coffee farms (in grey). Data are mean values ± SE, and the black dotted line represents the overall mean values.

Our results indicate that brood production and mortality varied significantly over time, but there was no significant difference between conventional and control groups. This suggests that pesticide exposure did not have a statistically significant impact on colony reproductive capacity and brood mortality. The significant variation over time may be associated with seasonal influences or other environmental factors. Differences in average brood mortality rates in different periods may result from natural fluctuations in colony population, as these managed colonies were recovering from the winter season and were transported to the farms and back to the forest patch, changing the ecological conditions in each transportation event, which can be a significant source of stress. Abrupt changes in the environmental and/or weather conditions, confinement and disturbance during transportation can increase brood mortality, impair brood production, foraging activity, and even the overall colony health. Furthermore, the rate of brood mortality in managed colonies placed on conventionally managed and chemical pesticide-free control coffee farms remained within the typical range for stingless bees^29^. In contrast, foraging activity showed significant differences between the conventional and control groups. In both groups, foraging activity was minimal before coffee blooming, but it was notably lower at conventionally managed farms. At this period, foragers had not yet been exposed to the nectar and pollen of coffee flowers. Therefore, other factors contributed to the difference on foraging activity, such as transportation stress, difficulty for foragers locating their nests in the crop field setting or other characteristic in the farming systems. However, in the first assessment after coffee bloom, foraging activity in both groups increased similarly, and colonies with low initial foraging rates recovered in the following weeks, equalizing between the two groups in subsequent assessments. For these reasons, the exposure to thiamethoxam should not be the primary cause of the difference in foraging activity.

Thiamethoxam and its metabolite clothianidin were detected in all coffee leaves sampled at conventional coffee-producing farms (Supplementary Table 2). Also, thiamethoxam residues were found in nectar and pollen stored in the stingless bee nests kept in all conventionally managed farms during coffee blooming (Supplementary Table 2). At the colony level for *A. mellifera*, effects on colony development were observed after 6-week continuous feeding with 100 ppb thiamethoxam, with no adverse effects at 50 ppb and no effects observed (NOEC) at 37.5 ppb thiamethoxam^45^. This NOEC for honeybees is 1.8 to 2-fold higher than the residues detected in the pollen and nectar collected from food stores of the stingless bee nests in this study. Although laboratory studies are crucial for indication the potential risks of pesticide use for the environment and non-target organisms, studies under realistic field settings are more reflective of the impacts of pesticides on pollinators, but significantly more challenging due to the inherent difficulty of conducting controlled experiments in the field^46^. For social pollinators, colony-level studies, which integrate all sublethal effects on individuals, are important to validate regulatory decision-making in evaluating the potential risk of bee-toxic pesticide use^25^; effects on colony growth and survival cannot be fully assessed by testing individuals in the laboratory. In such studies, attention should be given to the peculiarities of the farming system and characteristics of the studied agricultural crop. In this sense, for systemic insecticides, the results found for one specific crop should not be directly extrapolated for another crop without clear criteria^27^.

Overall, similar findings have been found in other field studies^46^. While laboratory-based studies have demonstrated that neonicotinoids can have sublethal effects at individual-level on foraging behaviour, cognitive abilities, and brood development, these detrimental effects have often not been observed in the field^46–48^. Here, we explore the primary hypothesis aimed at elucidating these apparent discrepancies. Exposure to sublethal doses can vary in duration, depending on the crop blooming period. In the case of mass-flowering arabica coffee, exposure typically occurs over a span of just 3-4 days. Additionally, the actual dose of neonicotinoids to which a bee colony is exposed may be much lower than the residue levels in stored pollen and nectar within the nest. Stingless bees visit a wide array of flowering plants to collect food resources^29^, which leads to a mixing of pollen and nectar from treated crops with pesticide-free plants, thereby diluting the residue levels. Stored pollen and nectar undergo multiple processing, storage, and exposure routes. Besides that, in general, systemic pesticide residues on flowers resulting from applications done much before blooming typically remain very low. In our study areas, applications were conducted an average of 8 months before coffee blooming, accounting for the low residue levels we detected. In apple orchards, for instance, no residues of systemic pesticides were detected in the whole flower, pollen and nectar sampled in the spring when applied via foliar spray in the previous fall^49^. Moreover, both the nutritional value of crops and wildflowers and the recovery mechanisms of social bees may compensate for negative pesticide effects^47,50^, allowing the costs of losing foraging workforce without compromising colony survival^27^. Further studies are necessary to explore the fate of these neonicotinoids within the colony, their actual exposure levels in stingless bee brood, adult workers, and the egg-laying queen.

The originality of our study lies in the combination of (1) manipulation of pollinator abundance in coffee producing-farms through the introduction of healthy, managed bee colonies to assess coffee yield, (2) the use of a comprehensive evaluation composed of a large number of physical, chemical and sensory parameters influencing coffee quality and market value (3) the use of managed stingless bees in higher-tier assessment to evaluate colony health under field-realistic exposure levels of the neonicotinoid thiamethoxam. Together, our results provide important insights into the combined use of managed and wild pollinators and enhancement of agroecosystem quality through pesticide stewardship and habitat management^21,22^. Thus, the deployment of managed bee colonies provides farmers with immediate means to raise coffee yield and beverage quality, mainly specialty coffee that increases farm-gate prices, alongside long-term sustainable profit to implement native forest restoration within coffee farms^43^. We highlight that pollination is a key input into intensively managed coffee-producing farms, and as agricultural production and environmental conservation interact synergically to both maximize profitability, which is fundamental to encourage farmers to be good stewards of their croplands by adopting nature-positive practices, and provide multi-functional benefits to society.

## Methods

### Study species

As highly eusocial insects, honeybees and stingless bees share many life-history traits^29,51^. Their colonies are swarm-founded, composed of one egg-laying queen that is morphologically distinct from the workers. With a complex chemical communication system and sophisticated division of labour based on age, older workers perform riskier tasks, such as foraging. As their perennial colonies require a constant provision of food throughout the year, generalist foragers collect nectar and pollen from a wide range of flowering plant species and store them within their nests. Nonetheless, while honeybees and stingless bees share these common traits of colony-forming insects, they differ in several lifestyle aspects and here we focused on biological traits from our study species.

The Africanised honeybees are a result of the admixture between African and European subspecies in Brazil since the 1950s that spread rapidly over the Americas in a few decades^52^. However, genetic and behavioural traits from the African subspecies (*Apis mellifera scutellata*) have been preserved during its widespread expansion^53^ enabling colonies to perform much better in tropical environments than their temperate-evolved relatives^41^. Compared to European subspecies, the Africanised honeybees are more productive, more defensive, rear proportionally more brood, have faster developmental time from egg to adults, and the shorter-lived workers are smaller and forage at younger age^52^. Also, as they exhibit higher growth rates in worker population, they produce more swarms annually as well as abandon their nests (i.e., abscond) much more readily when exposed to bad weather conditions, floral resource scarcity, and intense beekeeper handling^52^.

The stingless bee *Scaptotrigona depilis* uses tree cavities to establish their nests, which are built with a mixture of wax and plant resins (i.e., cerumen) that inhibit the growth of multiple bacteria and fungi due to their antimicrobial properties^41^. Nests are essentially composed of an entrance tube, egg-shaped pots to store pollen and honey and multilayered horizontal brood comb with same-sized cells to rear workers and males, while queens are reared in larger royal cells^49^. All brood cells are constructed, mass-provisioned with liquid larval food immediately before the queen lays her egg on top of it, and then sealed by workers^49^. In addition to larval food, *S. depilis* larvae consume *Zygosaccharomyces* fungus that grows inside brood cells and provides steroid precursors necessary for brood survival and metamorphosis^28^. The colonies typically contain around 10,000 adult workers headed by a mother queen who lays around 300 eggs daily^54^.

### Study sites

Our fieldwork was carried out in multiple coffee farms in the states of São Paulo and Minas Gerais in Southeastern Brazil (Extended Data Fig. 1), between 2021 and 2023 (Supplementary Table 1). This is the most traditional region for arabica coffee production in the country, with a large number of intensively managed conventional farms, relying on the use of chemical pesticides, and certified organic farms^30^. In all selected farms, arabica coffee was grown in full sun.

### Contributions of managed bees to coffee yield and quality

#### Managed bees and coffee production

One month before the coffee flowering season, we carefully selected healthy bee colonies according to the presence of an active egg-laying queen, the number of combs with brood, food stores, and worker population size. Five days before the onset of the first blooming season, between September and October 2021, we introduced colonies of the Africanised honeybee *A. mellifera* into 17 conventional farms (CA1–CA18; Supplementary Table 1), at a density of four colonies per hectare. At least 15 days before the second blooming season (2022), we supplemented six conventional farms with managed colonies of the stingless bee *S. depilis* (CS1– CS6; Supplementary Table 1) at ten colonies per hectare. For each farm, we assessed coffee yield at two distinct treatment areas: (1) at a distance of < 50 m from the colonies (‘with managed bees’) and (2) a more distant area from the bee colonies (mean distance: 250 m; range: 200–300 m, ‘without managed bees’)^33^.

Ripe coffee berries were manually or mechanically harvested between June and July 2022 in farms supplemented with honeybees (Supplementary Table 1). The manual method selected 1 to 3 sites in each treatment area, with and without managed bees. In each site, 7 to 100 coffee bushes were completely harvested, and the volume of ripe berries per bush was measured. The mechanical harvesting method consisted of harvesting ≥ 400 coffee bushes, measuring the volume of harvested berries and counting the number of bushes to estimate the volume of coffee berries per bush (in litres). Information of bushes per hectare provided by farmers was then used to calculate coffee yield per bush into 60 kg bags per hectare, as described by Almeida-Dias et al.^33^. On six farms supplemented with stingless bees, coffee berries were manually harvested between June and August 2023. In each site, we stablished a 10-m transect from stingless bee colonies, and we chose 10 branches at chest level from 10 different coffee bushes, totalling 360 branches (10 bushes × 3 sites × 2 treatment areas × 6 farms). Ripe berries were then collected, counted, and weighed to calculate the coffee yield (kg branch^-1^) for each site.

#### Managed honeybees and coffee quality

We selected **7** out of the 17 conventional farms where managed colonies of honeybees were introduced (CA1–CA7; Supplementary Table 1) and coffee berries were harvested, we evaluated coffee quality based on physical, chemical and sensory analyses (see Supplementary Information for detailed methods).

*a) Physical analyses.* The physical parameters assessed were raw bean size, with a series of sieves from #18 to 14, and the percentage of physical defects (i.e., misshapen berries with one seed (peaberries) and grading).
*b) Chemical composition analyses.* All chemical analyses were performed by the Agricultural Product Processing Research Centre (Federal University of Lavras, Lavras, Minas Gerais State). The chemical composition was quantified in raw (for sugars and organic acids) and roasted coffee beans (antioxidant activity, polyphenols, trigonelline, chlorogenic acid, caffeine, fatty acids and volatile compounds).
*c) Sensory analyses.* For sensory analyses of coffee brew, 10 attributes (acidity, aftertaste, balance, body, clean up, flavour, fragrance, sweetness, uniformity and overall impression) were evaluated by certified Specialty Coffee Association (SCA) Q-graders from Nucoffee (São Paulo, São Paulo State) to generate a final score for the beverage classification, a perception of the brew quality^40^. On the SCA 100-point scale^55^, cup scores above 80 are classified as specialty coffee (80–84.9: very good; 85–89.9: excellent).
*d) Market value.* Nucoffee specialists estimated the coffee market value for 60 kg bags based on coffee score and bean size (ungraded and ≥16-sieved beans, the latter are exportable beans), quoted on the New York Stock Exchange. All monetary values were gathered in Reais (R$; Brazilian currency) and converted into US dollars (US$) using the exchange rate of 4.8999 R$/US$ from 06 November of 2023 (Brazilian Central Bank).

### Effects of thiamethoxam-based products on bee colony health

We selected six conventionally (CS1–CS6; Supplementary Table 1) and two organically (OS1–OS2) managed coffee-producing farms, the latter as a chemical pesticide-free control. Based on the pest control efficiency (i.e., agronomic efficiency) in conventional arabica coffee farms, the commercially available thiamethoxam-based products Verdadero 600 WG and Actara 250 WG were applied, by soil drenching, during the berry expansion stage and pre-blooming period, respectively (Supplementary Table 3). Both systemic pesticides are commonly used to control some key coffee pests, such as the coffee leaf miner *Leucoptera coffeella*, one of the major threats to coffee production, and the fungus *Hemileia vastatrix*, a pathogen that causes the devastating disease coffee leaf rust.

To evaluate the effect of thiamethoxam and its metabolite clothianidin on stingless bee colony health, we introduced *S. depilis* colonies 10–18 days before coffee blooming between September and October 2022 (5 and 10 colonies on each conventional and organic farm, respectively). Among the temporal dynamics components of a bee colony, we assessed three colony performance traits: (1) brood production, (2) brood mortality rate, and (3) foraging activity. To assess (1) brood production, we counted the number of brood cells being built at the inspection moment – i.e., in the pre-provisioning and oviposition stage (for details, see Grüter^29^); (2) brood mortality rate, we recorded the number of empty brood cells and the total number of brood cells in a brood comb at pupal stage – i.e., percentage of dead brood removed from their cells by workers; (3) foraging activity, we counted the number of foragers leaving their colonies within a 3-min period from 09:00 to 12:00h. These parameters were recorded in five periods: 5–7 days before coffee blooming (pre-b), 5–7 (post-b), 45, 75 and 105 days after blooming. While evaluations during both pre-b and post-b were conducted in the coffee farms, the others were carried out in a stingless bee apiary in a 30-ha forest patch at the Brazilian Agriculture Research Corporation (Embrapa-Environment; Jaguariúna, São Paulo State), located over 90 km south of the most Southern coffee farm, where they were kept before the experiments. In addition, thiamethoxam and clothianidin residues were analysed in coffee leaves and in flower resources (nectar and pollen) collected by stingless bees at conventional farms during the blooming period (see Supplementary Information for detailed methods).

### Statistical analyses

All statistical analyses and figures were performed using R version 4.3.2.

#### Contributions of managed bees to coffee yields

While coffee productivity of farms supplemented with honeybee colonies was assessed in bags ha^-1^, productivity on farms with stingless bees was evaluated in terms of kg branch^-1^ bush^-1^. To address the non-independence of experimental design, given the foraging ranges of honeybees (mean 1.32 km, ref. ^35^) and *Scaptotrigona* bees (mean 0.87 km, ref. ^36^), as well as accounting for repeated measures at the sampling sites, we employed a more robust statistical model. This model incorporated random effects to enhance its ability to account for underlying variability within studied subjects. As a result, a linear mixed-effects model (LMM) was constructed, using bags ha^-1^ or kg branch^-1^ as proxies for coffee yields (i.e., the response variable) and the pollination treatment (with or without managed bees) as the fixed effect (i.e., predictor variable). To account for the hierarchical structure of our experimental design, the random effects in the LMM were configured with both cross-sectional and nested components for both managed bee species. Nevertheless, for honeybees, the first random effect was defined as ‘sampling units’ (intercepts) | ‘harvest method’ / ‘coffee variety’ (slopes), and the second random effect was structured as ‘farm property’ (intercepts), ‘farm ID’/ ‘plots within farms’ (slopes). Yet, for stingless bees, the single random effect was arranged as ‘sampling units’ (intercepts), with ‘plots within farms’ (slopes) nested with ‘farm ID’. The structure of the random effects in both LMMs was intentionally designed with intercepts to establish baseline values for some farm characteristics, and slopes were incorporated to capture variations within specific characteristics, thereby augmenting the capacity of LMMs to effectively account for and distinguish between sources of data variability.

Before data analyses, we standardized the response variable using the R function *scale*, thereby transforming the response variable with µ = 0 ± 1 standard-deviation to facilitate comparison with coffee farms supplemented with stingless bee colonies, for which we used a different metric as the response variable. Therefore, we fitted two LMMs, being one of them with the response variable log-transformed. Both models underwent checks for normality and homogeneity of variance across groups using the *check_normality* and *check_heteroscedasticity* functions, respectively, in *performance* R package^56^. Since only the latter log-transformed model showed adherence to normality (honeybees: *P* = 0.39; stingless bees: *P* = 0.48) and homogeneity (honeybees: *P* = 0.74; stingless bees: *P* = 0.73), they were chosen for further analysis. LMMs were performed using *lmer* function from *lme4* package^57^.

To understand the potential effects of pollination mediated by managing both bee species on coffee yields, we checked for the parameters and dispersion measures of the fixed effects of the best model fitted with restricted maximum likelihood (REML). To enhance the precision and efficiency of parameter estimation in LMM, we used Bound Optimization BY Quadratic Approximation (BOBYQA algorithm) as a method for bounded optimization through quadratic approximation. Finally, to assess the potential overdispersion in our LMM, we simulated residuals using the *simulateResiduals* function from *DHARMa* package^58^ to compare observed patterns in the raw data with patterns expected under the correct model specification. These results were visually analysed, checking for unusual patterns, heteroscedasticity, or any indications of overdispersion. All LMMs supported the assumptions above.

#### Contribution of managed honeybees to coffee quality

To evaluate the effects of pollination mediated by managed honeybees on coffee quality, we selected 42 coffee parameters comprised of seven major categories: physical parameters (*n* = 6; four sieve sizes (from #18 to 14), percentage of peaberries and grading), chemical composition in raw coffee beans (*n* = 5; antioxidant activity, polyphenols, and the bioactive compound trigonelline, chlorogenic acid, and caffeine), fatty acids (*n* = 4; palmitic, stearic, linoleic and araquidic acid), volatile compounds (*n* = 18; 2-methylfuran, butanal, 2-methylbutanal, 3-methylbutanal, imidazole, 2,3-pentanedione, 1-methylpyrrole, pyridine, pyrazine, methylpyrazine, 2-propanethiol, 2,5-dimethylpyrazine, 2,6-dimethylpyrazine, ethylpyrazine, furfural, furfuryl formate, 5-methyl-2-furancarboxaldehyde and 2-furanmethanol), sugars (*n* = 4; raffinose, sucrose, glucose, and fructose), organic acids (*n* = 4; citric, malic, quinic and succinic acid), brew quality (*n* = 1; coffee score). Furthermore, we evaluated the market value of 60 kg bag for ungraded and ≥16-sieved coffee beans.

To compare coffee quality from farms with and without managed honeybees, we performed a Procrustes Analysis (PA). Before this analysis, we standardized subsets (i.e., pollination treatment) using the Hellinger method and subsequently transformed them into two distance matrices with the Euclidean method. For this, we used the *decostand* and *vegdist* functions^59^. We executed PA with the *procrustes* function, and to test the congruence between the matrices, a permutation test (1,999 permutations) was performed using the *protest* function of *vegan* R package^59^.

The PA, a least-squares orthogonal mapping used to compare two multivariate datasets^60^, normalizes and rotates the ordination to achieve an optimal superimposition that maximizes the fit^60,61^. In this analysis, the degree of concordance, quantified as M^2^ (ref. ^60^), is determined by calculating the sum of squared residuals between configurations in their optimal superimposition. The M^2^ value ranges from 0 to 1, with lower values indicating a greater concordance among configurations^61^. Consequently, if both subsets exhibit similarity, the points in the rotated configuration should be closely grouped within the same subspace. By providing the trajectories and magnitudes of the residuals, PA offers valuable insights into the distinctive characteristics and variations observed in their coffee parameters in relation to pollination treatment in each of the seven farms. For instance, the positioning of residuals provides information on where corresponding points diverge^62–64^. Additionally, the displacement (trajectories) of residuals offers a dynamic perspective, illustrating the directional paths along which points undergo shifts and patterns of misalignment^62–64^. Finally, the variance (size) of residuals serves as a quantitative measure, indicating the overall magnitude of differences between the configurations: larger residuals suggest substantial dissimilarity, while smaller residuals signify closer agreement^62–64^.

Out of the 44 aforementioned coffee parameters, we then focused on the coffee score, market value of exportable beans (≥16-sieved beans; US$ bags^-1^), coffee bean size (passing sieve ≥16; %); peaberries (%), and caffeine content (g 100g^-1^). We chose these five parameters in line with most farmers who aim to meet prime quality parameters essential for satisfying specialty coffee market demands and ensuring exportation. These variables were tested in relation to pollination treatment across seven coffee-producing farms. Then, we fitted five generalized linear models (GLMs), with each of the five parameters serving as dependent variables. Pollination treatment was categorized by a fixed predictor variable, distinguishing between farms with and without managed honeybees, using Gamma family distribution and R-function *glm*. The assumptions for normality were evaluated using the *cvm.test* function in *goftest* R package^65^, while the homogeneity of variances was assessed by the *bptest* function from *lmtest* package^66^. All GLMs supported the aforementioned assumptions.

#### Effects of thiamethoxam-based products on bee colony health

To evaluate the effects of thiamethoxam-based products on bee colony health, we compared performance traits of *S. depilis* colonies introduced on conventional and organic managed coffee farms, using three generalized linear mixed models (GLMMs), where the response variables were brood production, brood mortality (%), and foraging activity, respectively. The predictor variables were the periods (pre- and post-blooming, and the subsequent 45, 75, and 105 days), the farming system (conventional *vs*. organic), and their interaction. As each group of managed colonies was introduced on different farms, we structured farms and colony IDs as crossed random effects. Model selection procedures were carried out before performing each of three GLMMs. First, we used the function *glmmTMB* from *glmmTMB* package^67^ to build independent models, using three error distributions: Poisson, quasi-Poisson, and negative binomial. Then, we selected the best model with the lowest Akaike information criterion (AIC), using the *AICtab* function from *bbmle* R package^68^. The parameters of GLMMs were extracted with the *Anova* function in *car* R package^69^. In addition, we used the *emmeans* function in *emmeans* package^70^ to extract estimated marginal means (EMMs) and their standard errors (± SE). To account for multiple comparisons, we adjusted the *P*-values using the false discovery rate (fdr) method when performing pairwise comparisons of grouping variables.

## Data availability

Data are available at Zenodo: https://doi.org/10.5281/zenodo.10975249

## Code availability

R scripts are all available to editors and referees upon request.

## Inclusion and ethics statement

The authorship is representative in terms of gender, scientific expertise, career stage and career paths, encompassing researchers from public institutions and private companies in both the Global South and North. The significance of this study extends beyond local relevance to a global level. No ethic approval was required. This study was carried out under beekeeping and transportation permits GEFAU 3810010/2021 from the Secretariat of Environment, Infrastructure and Logistics (Secretaria de Meio Ambiente, Infraestrutura e Logística) and GEDAVE 1639 from Coordination of Agricultural Defence (Coordenadoria de Defesa Agropecuária do Estado de São Paulo). Local beekeepers held their own beekeeping permits for colony transportation.

## Acknowledgements

We are grateful to farmers and beekeepers for their support to this study, allowing us to access coffee farms, assisting with coffee harvest and bee management. We also thank the Embrapa teamwork for helping with field logistics. This work was supported by Embrapa and Syngenta (grant SEG 10.20.00.143.00.00), AgroBee Pollination and Sustainability Solutions Startup, the National Council for Scientific and Technological (CNPq, grants 164743/2020-0 to D.A.A. and 350679/2022-3 to C.F.S.), the São Paulo Research Foundation (FAPESP, grants 2018/23805-8, 2017/07848-6, 2020/03549-7, to D.M.O., A.A.B., G.J.G.S., respectively), and the Studies and Projects Funding Agency (FINEP, grant 17/50362-7 to J.M.V.A.D. and A.A.B.).

## Author contributions

Conceptualization: J.D.R., D.M.O., J.M.V.A.D., A.P.C., A.J.C., A.A.B., C.M.; Methodology: J.D.R., D.M.O., J.M.V.A.D., A.P.C., A.J.C., A.M.P., A.C.M.Q., C.M.; Investigation: J.D.R., D.M.O., J.M.V.A.D., F.C.L.Q.; Data curation: D.A.A., J.D.R., C.F.S., D.M.O.; Formal analysis: C.F.S.; Validation: C.F.S.; Supervision: A.A.B., J.M.S.B., C.M.; Project administration: A.P.C., N.B.T.V., A.C.M.Q., C.M.; Resources: D.A.A., D.M.O., J.M.V.A.D., G.S.S., T.S.O.K., A.P.C., A.A.B., G.J.G.S., C.M.; Writing – original draft: D.A.A., J.D.R. All authors contributed to reviewing and editing the final manuscript.

---

**Extended Data Table 1.**
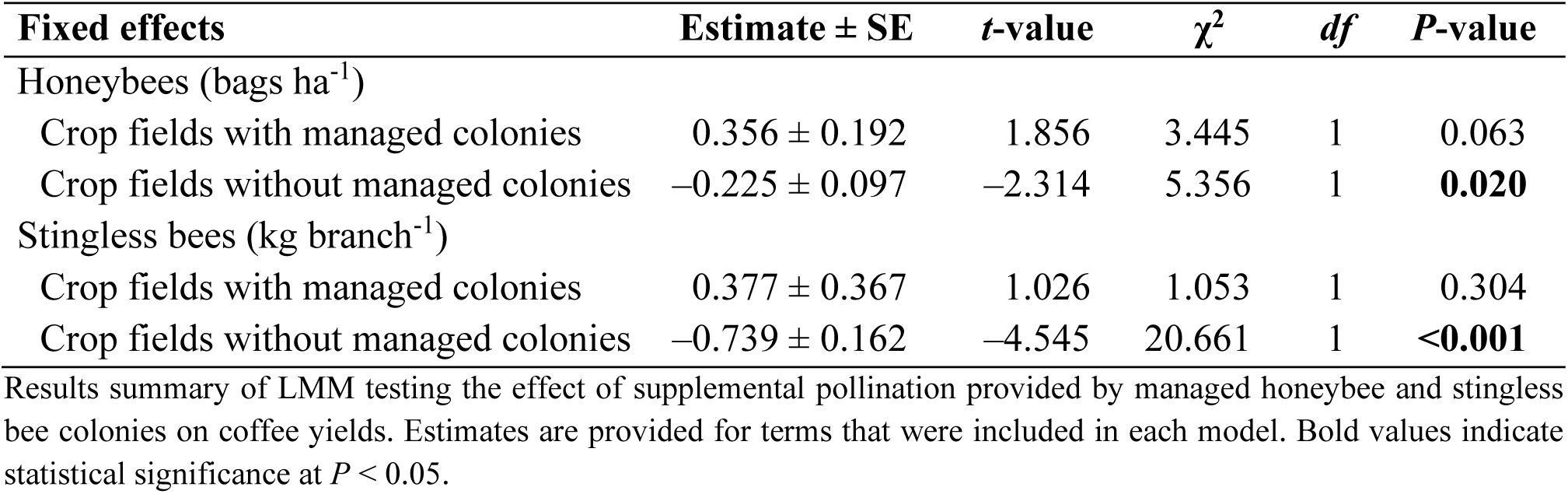
| Effect of managed bee colonies on arabica coffee yields.

**Extended Data Fig. 1.**
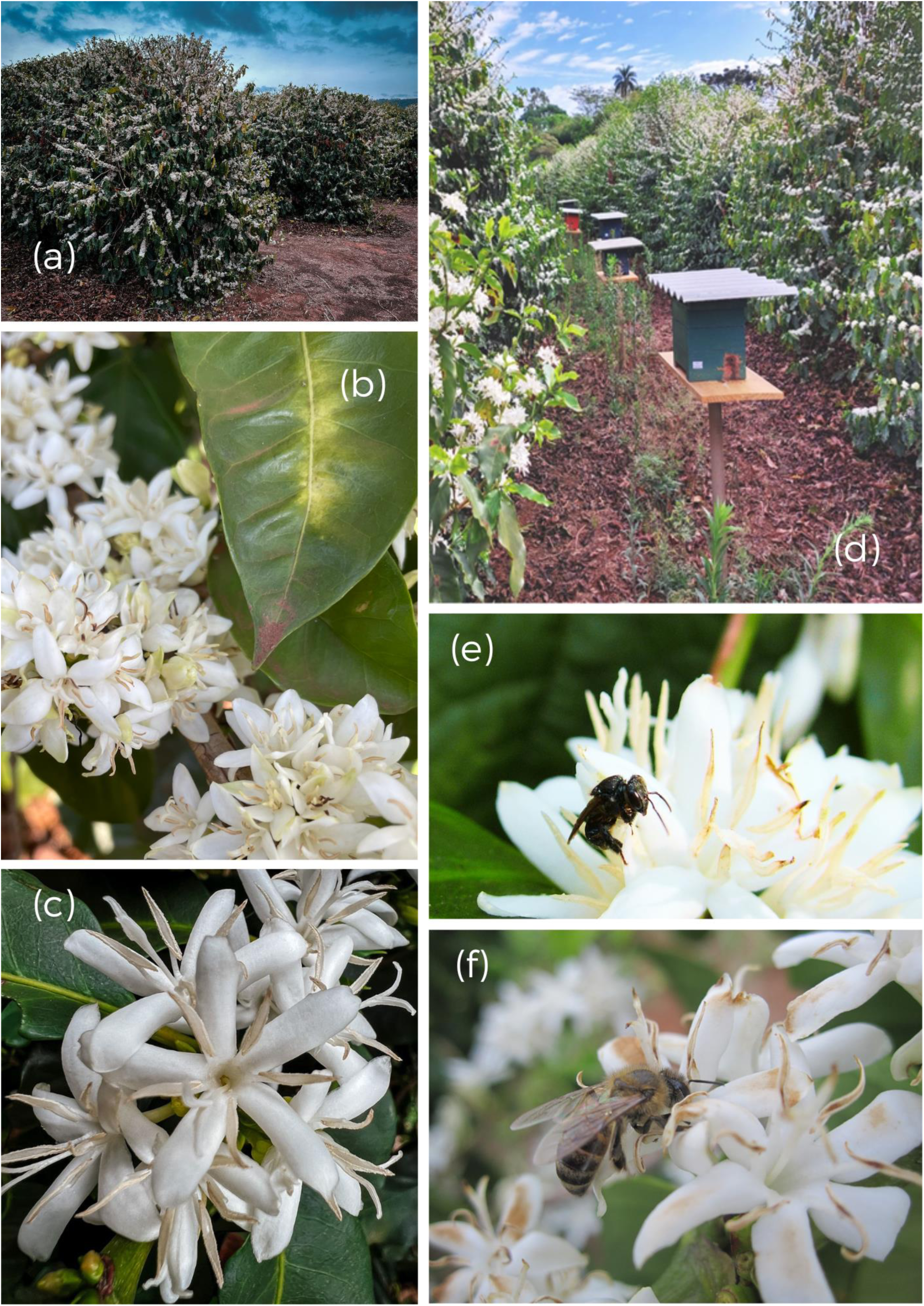
| Conventional arabica coffee-producing farms supplemented with managed bee colonies. Mass-blooming coffee bushes (a) that boast numerous inflorescences (b), each grouping hermaphrodite flowers (c), which only open for 3–4 days. Managed bee colonies, kept in wooden hives, introduced into coffee field (d) to ensure supplemental pollination provided by stingless bees (e) and honeybees (f).

