## SUPPLEMENTARY INFORMATION for "A sustainable and bee-pollinated coffee to start your day"

### Contents

Supplementary Methods

Supplementary References

Supplementary Tables 1–5

### Supplementary Methods

#### 1. Coffee quality

#### 1.1. Chemical composition analyses

For the chemical analyses, the samples were prepared by grounding raw and roasted coffee beans in an 11A basic mill (IKA, Brazil) for approximately 1 min, with the addition of liquid nitrogen to facilitate grinding and prevent oxidation. The ground samples were packed in falcon tubes and stored at –85 °C until the analyses were conducted.

**(a) Organic acids**

For the extraction of organic acids, 0.25 g of ground raw coffee was weighed and placed in falcon tubes with 25 mL of 0.2% perchloric acid. The solution was stirred for 10 min. Subsequently, the solution was filtered through common filter paper and then filtered through a 0.45 µm membrane. The determination of these compounds was done by High-Performance Liquid Chromatography (HPLC) based on the methodology described by Jham et al.[1], using 0.2% perchloric acid as the mobile phase, with a flow rate of 1 $mL.min^{-1}$ and the C610H column at 40 °C, monitored by UV spectrophotometry at 210 and 254 $\times 10^{-9}$ m wavelengths. Standard solutions of citric, malic, tartaric, succinic, lactic, quinic, and acetic acids were used for peak identification in the chromatogram, comparison of retention times, and for calculating their concentration in the samples. The final levels of organic acids were given in mg/100 gd.m. and mg/100 gd.m..

**(b) Sugar profile**

For sugar extraction, 100 mg of ground and degreased raw coffee samples contained in 2 mL microcentrifuge tubes were suspended in 1 mL of ultrapure water. The tubes were placed in a heated ultrasonic bath at 80 °C for 15 min. Then, the samples were centrifuged for 3 min at 5,500 rpm. An aliquot of 500 µL of the extract was transferred to another 1.5 mL microcentrifuge tube and then centrifuged for 5 min at 5,500 rpm. The supernatant was filtered through a 0.45 μm membrane filter and directly injected into an HPLC. The determination of these compounds was performed using an Agilent chromatograph with a refractive index detection system, using a Supelco 610H chromatographic column. The mobile phase consisted of 1% phosphoric acid-acidified water (at a flow rate of 0.5 $mL.min^{-1}$). For identification and quantitative analysis, a standard curve was prepared using raffinose, sucrose, glucose, fructose, and arabinose standards. The final content of sugars was given in g of sugar per 100 g of dry matter (g/100 gd.m.).

**(c) Antioxidant activity**

The antioxidant activity by the DPPH method was determined according to Rufino et al.[2] using 1 g of ground roasted coffee to 4 mL of water at different dilutions. The absorbances were measured using UV-VIS at 515 nm.

**(d) Polyphenols**

Phenolic compounds were extracted at a ratio of 1 g of roasted coffee to 4 mL of water. The polyphenols were assayed using the Folin-Denis reagent at 760 nm[3]. The results were expressed in mg of gallic acid per 100 g of sample.

**(e) Bioactive compounds**

The non-volatile compounds caffeine, trigonelline, and chlorogenic acids were determined by HPLC (methodology adapted from Vitorino et al.[4]). Samples of 0.25 g of ground raw coffee were extracted in 25 mL boiling distilled water and placed in a water bath, with boiling water, for 3 min. The extract was filtered through common filter paper and then filtered through a 0.45 μm membrane. The determination of these compounds was performed using an Agilent chromatograph with ultraviolet detection system, a C18 chromatographic column, and a wavelength of 272 nm. The mobile phase consisted of 85% A (1% acetic acid aqueous solution) and 15% B (methanol), at a flow rate of 1 mL min$^{-1}$. For identification and quantitative analysis, a standard curve was prepared using caffeine, trigonelline, and 5-caffeoylquinic acid (5-CQA) standards.

**(f) Fatty acids**

To determine fatty acids, 250 mg of ground green coffee beans from each sample were weighed and placed in 2.5 mL centrifuge tubes, to which 1.0 mL of hexane was added. The tubes were then placed and kept in an ultrasonic bath for 10 min at room temperature. Subsequently, the tubes were centrifuged for 2 min at 5,000 rpm. A portion of 500 µL of each supernatant was transferred to 2.0 mL cryogenic tubes. The residual hexane in the aliquot was evaporated in the fume hood, and the fatty acid content was subjected to lipid hydrolysis steps. For lipid hydrolysis, approximately 10 mg of extracted oil was diluted in 100 µL of ethanol (95%) and 1 mol L$^{-1}$ potassium hydroxide solution (5%). After 10 s of vortex agitation, the oil was hydrolysed using a conventional 80 W Panasonic® microwave oven for 5 min. After cooling, 400 µL of 20% hydrochloric acid, a small amount of NaCl, and 600 µL of ethyl acetate were added. After 10 s of vortex agitation and 5 min of rest, 300 µL of the organic layer was removed, placed in microcentrifuge tubes, and dried by evaporation, obtaining the free fatty acids (adapted from Christie et al.[5]).

The free fatty acids were methylated with 100 µL of 14% methanol BF3 and heated in a water bath at 80 °C for 10 min. They were then diluted with 600 µL of hexane and analysed by gas chromatography. From each extract, 2 µL were injected into a gas chromatograph (GC-2010, Shimadzu) coupled with a mass spectrometer detector, equipped with a SP-2560 column (Supelco) 100 m × 0.25 mm, was used with a temperature gradient of 140 °C for 5 min, 4 °C min$^{-1}$ up to 240 °C, remaining at this temperature for 30 min; Injector (1/20 split), at 240 °C, and detector at 240 °C. Helium was used as carrier gas with a linear velocity at 2 mL min$^{-1}$. The peaks corresponding to each fatty acid were identified by comparing them with Supelco37 methylated fatty acid standards. The final levels were given as a percentage of the relative area.

**(g) Volatile compounds**

The volatile compounds were analysed using gas chromatography of ground samples from the roasted beans used in the sensory analysis. After grinding, 1 g of each sample was placed in hermetically sealed vials. The volatile compounds were extracted using the static headspace of the gas chromatograph coupled with a mass spectrometer (GC-MS; QP-2010 SE; Shimadzu, Japan), equipped with an NST-100 column (30 m × 0.25 mm × 0.25 μm) with a polyethylene glycol phase similar to Carbowax®. The vials containing the samples were placed in the equipment. After equilibration at 70 °C for 30 min, the volatile phase was injected into the gas chromatograph with subsequent detection using a MS. The injector temperature was set at 220 °C, and helium gas was used as a carrier and maintained at a linear velocity of 1 mL $min^{-1}$. The heating program was as follows: for 6 min, the oven temperature was maintained at 25 °C, then increased to 70 °C at a rate of 10 °C $min^{-1}$, up to 95 °C at 5 °C $min^{-1}$, up to 115 °C at 10 °C $min^{-1}$, up to 170 °C at 5 °C $min^{-1}$, and finally up to 215 °C at 40 °C $min^{-1}$. The total run time was 35 min.

The data analysis and the compounds' identification were conducted using the GCMS Solution software (version 4.4) and the NIST/EPA/NIH 2014 database. Chemical identification was performed by comparing the MS spectra with the database. The results were expressed as a relative percentage area corresponding to the peak area of each identified compound, calculated from the total area of the chromatogram.

**1.2. Sensory analyses of brewed coffee**

The coffee beans were dried until their moisture content reached 11%, transferred to storage bins for a minimum of a five days and hulled for removal of the parchment. Then, the raw coffee beans were roasted until a medium roasting level in the scale of Specialty Coffee Association (SCA[6]) and after 8-10 hours, the roasted beans were ground.

Following SCA protocol[6], the evaluation of the coffee quality was performed by Q-graders. First, roasted samples were visually inspected for colour and then five transparent cups were prepared for each sample containing 8.5 g of ground coffee and covered. The fragrance was evaluated by the tasters 15 min after grinding. Then, water at 93-94 °C was added over the ground coffee, and fragrance was evaluated after 3-5 min rest. The other attributes (acidity, aftertaste, balance, body, clean up, flavour, sweetness, uniformity and overall impression) were individually evaluated after 8-10 min. Finally, for each sample, tasters assigned a global score, which corresponds to the brew quality (> 80.0: not specialty; 80.0-84.9: very good specialty, 85.0-89.9: excellent specialty; > 90.0: outstanding specialty).

**2. Pesticide residue analyses**

Thiamethoxam and clothianidin residues were analysed in coffee leaves (at least 10 leaves from different bushes were sampled per farm) and in flower resources collected by *S. depilis* foragers at six conventional farms during blooming period (CS1–CS6; Supplementary Table 1). To assess them, in late afternoon, we sampled recently collected pollen (mean ± SD = 0.502 ± 0.219 g/sample, $n$ = 17 pollen samples) nectar (1.578 ± 0.689 g/sample, $n$ = 17 nectar samples) directly from open food storage pots within 3–5 stingless bee nests for each farm, as described in Menezes et al.[7]. All samples were stored at –2 °C for pesticide residue analyses performed at Eurofins Agroscience Services (Indaiatuba, São Paulo State).

To screen for thiamethoxam and its metabolite clothianidin, each pollen (50 ± 10 mg sample$^{-1}$ farm$^{-1}$) and nectar sample (150 mg sample$^{-1}$ farm$^{-1}$) were placed in a centrifuge tube. Water, acetonitrile and a mixture of salts were added and then vortexed. For pollen, after centrifugation, the supernatant was transferred to a vial and frozen, and a portion of extract was transferred to a vial with combination of salts and C18 for the clean-up step, followed by evaporation and resuspension in acetonitrile. For nectar samples, after centrifugation, a portion was firstly subjected to clean-up followed by partial evaporation and resuspension in acetonitrile. Finally, for determination of neonicotinoid residues in coffee bushes, 2.5 g of each leaf sample per farm were added to a centrifuge tube with water and acetonitrile and then vortex. Following centrifugation, a portion of extract was transferred to a vial containing mixture of salts for clean-up and activated charcoal, vortexed, centrifuged and filtered. For all three matrices, the final solution was analysed by liquid chromatography tandem mass spectrometry system (LC (Agilent)-MS/MS (SCIEX 5500)), using 0.05% acetic acid + ammonium formate in water as mobile phase A, and 0.05% acetic acid in methanol as mobile phase B, with a C18 column of 150 mm × 2mm × 5µm.

### Supplementary Tables

**Supplementary Table 1.** Coffee-producing farms supplemented with managed bee colonies (MG: Minas Gerais State; SP: São Paulo State).

| Farm | Farming system | Locality | Managed bee species | Dates when bee colonies were introduced on farms | Coffee harvesting date | Coffee harvesting method |
|---|---|---|---|---|---|---|
| CA1 | Conventional | Cambuquira, MG | *Apis mellifera* | October 2021 | June 2022 | Manual |
| CA2 | Conventional | Monte Carmelo, MG | *Apis mellifera* | October 2021 | June 2022 | Mechanical |
| CA3 | Conventional | Coromandel, MG | *Apis mellifera* | October 2021 | June 2022 | Mechanical |
| CA4 | Conventional | Santa Rosa da Serra, MG | *Apis mellifera* | October 2021 | June 2022 | Manual |
| CA5 | Conventional | Monte Carmelo, MG | *Apis mellifera* | October 2021 | June 2022 | Mechanical |
| CA6 | Conventional | Coromandel, MG | *Apis mellifera* | October 2021 | June 2022 | Manual |
| CA8 | Conventional | Carmo do Paranaíba, MG | *Apis mellifera* | September 2021 | June 2022 | Mechanical |
| CA9 | Conventional | Carmo do Paranaíba, MG | *Apis mellifera* | September 2021 | June 2022 | Manual |
| CA10 | Conventional | Patrocínio, MG | *Apis mellifera* | October 2021 | June 2022 | Mechanical |
| CA11 | Conventional | Patrocínio, MG | *Apis mellifera* | September 2021 | June 2022 | Mechanical |
| CA12 | Conventional | Coromandel, MG | *Apis mellifera* | October 2021 | June 2022 | Manual |
| CA13 | Conventional | Coromandel, MG | *Apis mellifera* | October 2021 | June 2022 | Manual |
| CA14 | Conventional | Pratinha, MG | *Apis mellifera* | September 2021 | July 2022 | Manual |
| CA15 | Conventional | Machado, MG | *Apis mellifera* | September 2021 | June 2022 | Manual |
| CA16 | Conventional | Altinópolis, SP | *Apis mellifera* | October 2021 | June 2022 | Mechanical |
| CA17 | Conventional | Patrocínio, MG | *Apis mellifera* | September 2021 | June 2022 | Mechanical |
| CA18 | Conventional | Alterosa, MG | *Apis mellifera* | September 2021 | June 2022 | Manual |
| CS1 | Conventional | Espírito Santo do Pinhal, SP | *Scaptotrigona depilis* | September 2022 | May 2023 | Manual |
| CS2 | Conventional | Altinópolis, SP | *Scaptotrigona depilis* | September 2022 | July 2023 | Manual |
| CS3 | Conventional | Franca, SP | *Scaptotrigona depilis* | September 2022 | July 2023 | Manual |
| CS4 | Conventional | São Sebastião do Paraíso, SP | *Scaptotrigona depilis* | September 2022 | July 2023 | Manual |
| CS5 | Conventional | São Tomás de Aquino, MG | *Scaptotrigona depilis* | September 2022 | June 2023 | Manual |
| CS6 | Conventional | Ibiraci, MG | *Scaptotrigona depilis* | September 2022 | July 2023 | Manual |
| OS1 | Organic | Franca, SP | *Scaptotrigona depilis* | September 2022 | June 2023 | - |
| OS2 | Organic | Franca, SP | *Scaptotrigona depilis* | September 2022 | June 2023 | - |

**Supplementary Table 2.** Field rates of thiamethoxam-based products applied by soil drenching on conventionally managed arabica coffee farms, date of applications, and residue levels (mean ± SD) of thiamethoxam and clothianidin detected in coffee leaves and in nectar and pollen stored within stingless bee nests sampled in September 2022.

| **Farm** | **Field rates (kg ha$^{-1}$)** [a,b] | | **Application dates** | | **Thiamethoxam (mg kg$^{-1}$)** | | | **Clothianidin (mg kg$^{-1}$)** | | |
|---|---|---|---|---|---|---|---|---|---|---|
| | Verdadero 600® WG [c] | Actara 250® WG [c] | Verdadero 600® WG | Actara 250® WG | Leaves | Nectar | Pollen | Leaves | Nectar | Pollen |
| CS1 | 1.00 | 1.00 | Nov 2021 | Feb 2022 | 0.1071 | 0.0052 ± 0.0010 | 0.0070 ± 0.0017 | 0.0509 | < 0.0010 | 0.0020 ± 0.0010 |
| CS2 | 1.00 | not applied | Nov 2021 | - | 0.0353 | 0.0034 ± 0.0001 | 0.0034 ± 0.0013 | 0.0349 | < 0.0010 | 0.0015 ± 0.0007 |
| CS3 | 1.00 | not applied | Nov 2021 | - | 0.0093 | 0.0026 ± 0.0001 | 0.0016 ± 0.0006 | 0.0102 | < 0.0010 | < 0.0010 |
| CS4 | 1.00 | not applied | Nov 2021 | - | 0.0317 | 0.0025 ± 0.0015 | 0.0037 ± 0.0014 | 0.0340 | < 0.0010 | < 0.0010 |
| CS5 | not applied | 1.00 | - | Dec 2021 | 0.0418 | 0.0214 ± 0.0014 | 0.0174 ± 0.0030 | 0.0327 | < 0.0010 | 0.0010 [d] |
| CS6 | 1.00 | 1.00 | Nov 2021 | Jan 2022 | 0.0110 | 0.0032 ± 0.0004 | 0.0045 ± 0.0006 | 0.0320 | < 0.0010 | < 0.0010 |

[a] Active ingredient: 30% and 25% thiamethoxam in formulated products Verdadero and Actara, respectively.
[b] Field rates based on the efficiency in coffee pest control.
[c] Label recommendations: 0.7–1.0 kg ha$^{-1}$ and 1.4–2.0 kg ha$^{-1}$ for Verdadero and Actara, respectively.
[d] Average value estimated for two pollen samples (<0.001 and 0.0020).

**Supplementary Table 3.** Pairwise comparisons of brood production by managed stingless bee colonies among experimental periods (pre- and post-blooming (b), and the subsequent 45, 75 and 105 days after blooming) and coffee farming systems (conventional and organic). Bold values indicate statistical significance at $P < 0.05$.

| Pair 1 | Pair 2 | Ratio | Std. Error | z-ratio | *P* -value |
|---|---|---|---|---|---|
| pre-b, organic | pre-b, conventional | 0.91 | 0.13 | –0.63 | 0.60 |
| pre-b, organic | post-b, conventional | 0.68 | 0.10 | –2.73 | **0.01** |
| pre-b, organic | 75 days, conventional | 0.66 | 0.09 | –3.02 | **0.01** |
| pre-b, organic | 45 days, conventional | 0.76 | 0.11 | –1.92 | 0.09 |
| pre-b, organic | 105 days, conventional | 0.49 | 0.07 | –5.24 | **<0.0001** |
| post-b, conventional | pre-b, conventional | 1.35 | 0.15 | 2.72 | **0.01** |
| post-b, organic | pre-b, conventional | 1.71 | 0.21 | 4.38 | **<0.0001** |
| post-b, organic | pre-b, organic | 1.87 | 0.24 | 4.80 | **<0.0001** |
| post-b, organic | post-b, conventional | 1.27 | 0.15 | 2.06 | 0.07 |
| post-b, organic | 75 days, conventional | 1.23 | 0.14 | 1.88 | 0.10 |
| post-b, organic | 45 days, conventional | 1.42 | 0.17 | 2.90 | **0.01** |
| post-b, organic | 105 days, conventional | 0.92 | 0.10 | –0.74 | 0.53 |
| 75 days, conventional | pre-b, conventional | 1.39 | 0.14 | 3.13 | **<0.0001** |
| 75 days, conventional | post-b, conventional | 1.03 | 0.10 | 0.31 | 0.79 |
| 75 days, organic | pre-b, conventional | 1.65 | 0.20 | 4.14 | **<0.0001** |
| 75 days, organic | pre-b, organic | 1.81 | 0.24 | 4.58 | **<0.0001** |
| 75 days, organic | post-b, conventional | 1.23 | 0.14 | 1.80 | 0.11 |
| 75 days, organic | post-b, organic | 0.97 | 0.10 | –0.32 | 0.79 |
| 75 days, organic | 75 days, conventional | 1.19 | 0.13 | 1.61 | 0.15 |
| 75 days, organic | 45 days, conventional | 1.37 | 0.16 | 2.65 | **0.02** |
| 75 days, organic | 105 days, conventional | 0.90 | 0.09 | –1.05 | 0.36 |
| 45 days, conventional | pre-b, conventional | 1.21 | 0.14 | 1.65 | 0.14 |
| 45 days, conventional | post-b, conventional | 0.90 | 0.10 | –1.03 | 0.36 |
| 45 days, conventional | 75 days, conventional | 0.87 | 0.09 | –1.37 | 0.22 |
| 45 days, organic | pre-b, conventional | 1.38 | 0.18 | 2.47 | **0.03** |
| 45 days, organic | pre-b, organic | 1.51 | 0.21 | 3.00 | **0.01** |
| 45 days, organic | post-b, conventional | 1.03 | 0.13 | 0.20 | 0.86 |
| 45 days, organic | post-b, organic | 0.81 | 0.09 | –1.93 | 0.09 |
| 45 days, organic | 75 days, conventional | 1.00 | 0.12 | –0.04 | 0.97 |
| 45 days, organic | 75 days, organic | 0.83 | 0.09 | –1.65 | 0.14 |
| 45 days, organic | 45 days, conventional | 1.14 | 0.15 | 1.05 | 0.36 |
| 45 days, organic | 105 days, conventional | 0.75 | 0.09 | –2.53 | **0.02** |
| 105 days, conventional | pre-b, conventional | 1.85 | 0.18 | 6.19 | **<0.0001** |
| 105 days, conventional | post-b, conventional | 1.37 | 0.12 | 3.52 | **<0.0001** |
| 105 days, conventional | 75 days, conventional | 1.33 | 0.11 | 3.43 | **<0.0001** |
| 105 days, conventional | 45 days, conventional | 1.53 | 0.15 | 4.44 | **<0.0001** |
| 105 days, organic | pre-b, conventional | 1.96 | 0.23 | 5.68 | **<0.0001** |
| 105 days, organic | pre-b, organic | 2.15 | 0.27 | 6.02 | **<0.0001** |
| 105 days, organic | post-b, conventional | 1.46 | 0.16 | 3.36 | **<0.0001** |
| 105 days, organic | post-b, organic | 1.15 | 0.11 | 1.42 | 0.21 |
| 105 days, organic | 75 days, conventional | 1.41 | 0.15 | 3.24 | **<0.0001** |
| 105 days, organic | 75 days, organic | 1.19 | 0.11 | 1.77 | 0.12 |
| 105 days, organic | 45 days, conventional | 1.62 | 0.19 | 4.18 | **<0.0001** |
| 105 days, organic | 45 days, organic | 1.42 | 0.15 | 3.29 | **<0.0001** |
| 105 days, organic | 105 days, conventional | 1.06 | 0.11 | 0.58 | 0.62 |

**Supplementary Table 4.** Pairwise comparisons of brood mortality by managed stingless bee colonies among experimental periods (pre- and post-blooming (b), and the subsequent 45, 75 and 105 days after blooming) and coffee farming systems (conventional and organic). Bold values indicate statistical significance at $P < 0.05$.

| Pair 1 | Pair 2 | Ratio | Std. Error | z-ratio | *P* -value |
|---|---|---|---|---|---|
| pre-b, organic | pre-b, conventional | 0.81 | 0.24 | –0.72 | 0.58 |
| pre-b, organic | post-b, conventional | 0.70 | 0.21 | –1.21 | 0.32 |
| pre-b, organic | 75 days, conventional | 3.57 | 1.17 | 3.90 | **<0.0001** |
| pre-b, organic | 45 days, conventional | 1.45 | 0.44 | 1.23 | 0.32 |
| pre-b, organic | 105 days, conventional | 1.85 | 0.59 | 1.94 | 0.09 |
| post-b, conventional | pre-b, conventional | 1.16 | 0.30 | 0.58 | 0.65 |
| post-b, organic | pre-b, conventional | 1.10 | 0.35 | 0.30 | 0.82 |
| post-b, organic | pre-b, organic | 1.36 | 0.46 | 0.91 | 0.48 |
| post-b, organic | post-b, conventional | 0.95 | 0.30 | –0.16 | 0.90 |
| post-b, organic | 75 days, conventional | 4.85 | 1.67 | 4.57 | **<0.0001** |
| post-b, organic | 45 days, conventional | 1.97 | 0.63 | 2.10 | 0.06 |
| post-b, organic | 105 days, conventional | 2.51 | 0.85 | 2.71 | **0.01** |
| 75 days, conventional | pre-b, conventional | 0.23 | 0.07 | –5.16 | **<0.0001** |
| 75 days, conventional | post-b, conventional | 0.20 | 0.06 | –5.68 | **<0.0001** |
| 75 days, organic | pre-b, conventional | 0.23 | 0.08 | –4.13 | **<0.0001** |
| 75 days, organic | pre-b, organic | 0.28 | 0.11 | –3.35 | **<0.0001** |
| 75 days, organic | post-b, conventional | 0.19 | 0.07 | –4.54 | **<0.0001** |
| 75 days, organic | post-b, organic | 0.21 | 0.08 | –3.99 | **<0.0001** |
| 75 days, organic | 75 days, conventional | 0.99 | 0.38 | –0.02 | 0.98 |
| 75 days, organic | 45 days, conventional | 0.40 | 0.15 | –2.49 | **0.03** |
| 75 days, organic | 105 days, conventional | 0.51 | 0.20 | –1.76 | 0.12 |
| 45 days, conventional | pre-b, conventional | 0.56 | 0.15 | –2.21 | 0.05 |
| 45 days, conventional | post-b, conventional | 0.48 | 0.12 | –2.85 | **0.01** |
| 45 days, conventional | 75 days, conventional | 2.46 | 0.72 | 3.09 | **0.01** |
| 45 days, organic | pre-b, conventional | 0.85 | 0.24 | –0.59 | 0.65 |
| 45 days, organic | pre-b, organic | 1.05 | 0.32 | 0.15 | 0.90 |
| 45 days, organic | post-b, conventional | 0.73 | 0.20 | –1.12 | 0.36 |
| 45 days, organic | post-b, organic | 0.77 | 0.25 | –0.80 | 0.53 |
| 45 days, organic | 75 days, conventional | 3.74 | 1.17 | 4.22 | **<0.0001** |
| 45 days, organic | 75 days, organic | 3.77 | 1.40 | 3.58 | **<0.0001** |
| 45 days, organic | 45 days, conventional | 1.52 | 0.44 | 1.45 | 0.22 |
| 45 days, organic | 105 days, conventional | 1.93 | 0.59 | 2.17 | 0.06 |
| 105 days, conventional | pre-b, conventional | 0.44 | 0.12 | –3.06 | **0.01** |
| 105 days, conventional | post-b, conventional | 0.38 | 0.11 | –3.45 | **<0.0001** |
| 105 days, conventional | 75 days, conventional | 1.93 | 0.60 | 2.12 | 0.06 |
| 105 days, conventional | 45 days, conventional | 0.79 | 0.23 | –0.85 | 0.51 |
| 105 days, organic | pre-b, conventional | 0.20 | 0.07 | –4.48 | **<0.0001** |
| 105 days, organic | pre-b, organic | 0.24 | 0.09 | –3.69 | **<0.0001** |
| 105 days, organic | post-b, conventional | 0.17 | 0.06 | –4.93 | **<0.0001** |
| 105 days, organic | post-b, organic | 0.18 | 0.07 | –4.31 | **<0.0001** |
| 105 days, organic | 75 days, conventional | 0.87 | 0.34 | –0.36 | 0.81 |
| 105 days, organic | 75 days, organic | 0.88 | 0.38 | –0.31 | 0.82 |
| 105 days, organic | 45 days, conventional | 0.35 | 0.13 | –2.86 | **0.01** |
| 105 days, organic | 45 days, organic | 0.23 | 0.09 | –3.95 | **<0.0001** |
| 105 days, organic | 105 days, conventional | 0.45 | 0.17 | –2.10 | 0.06 |

**Supplementary Table 5.** Pairwise comparisons of foraging activity by managed stingless bee colonies among experimental periods (pre- and post-blooming (b), and the subsequent 45, 75 and 105 days after blooming) and coffee farming systems (conventional and organic). Bold values indicate statistical significance at $P < 0.05$.

| Pair 1 | Pair 2 | Ratio | Std. Error | z-ratio | *P*-value |
|---|---|---|---|---|---|
| pre-b, organic | pre-b, conventional | 2.83 | 0.59 | 4.97 | **<0.0001** |
| pre-b, organic | post-b, conventional | 1.18 | 0.22 | 0.86 | 0.51 |
| pre-b, organic | 75 days, conventional | 0.90 | 0.16 | –0.59 | 0.66 |
| pre-b, organic | 45 days, conventional | 1.07 | 0.20 | 0.38 | 0.78 |
| pre-b, organic | 105 days, conventional | 0.76 | 0.14 | –1.56 | 0.19 |
| post-b, conventional | pre-b, conventional | 2.41 | 0.37 | 5.74 | **<0.0001** |
| post-b, organic | pre-b, conventional | 4.36 | 0.88 | 7.29 | **<0.0001** |
| post-b, organic | pre-b, organic | 1.54 | 0.19 | 3.41 | **<0.0001** |
| post-b, organic | post-b, conventional | 1.81 | 0.32 | 3.32 | **<0.0001** |
| post-b, organic | 75 days, conventional | 1.38 | 0.24 | 1.86 | 0.11 |
| post-b, organic | 45 days, conventional | 1.65 | 0.29 | 2.84 | **0.01** |
| post-b, organic | 105 days, conventional | 1.16 | 0.20 | 0.88 | 0.51 |
| 75 days, conventional | pre-b, conventional | 3.16 | 0.47 | 7.72 | **<0.0001** |
| 75 days, conventional | post-b, conventional | 1.31 | 0.15 | 2.36 | **0.04** |
| 75 days, organic | pre-b, conventional | 3.82 | 0.78 | 6.60 | **<0.0001** |
| 75 days, organic | pre-b, organic | 1.35 | 0.18 | 2.30 | **0.04** |
| 75 days, organic | post-b, conventional | 1.59 | 0.29 | 2.56 | **0.02** |
| 75 days, organic | post-b, organic | 0.88 | 0.10 | –1.12 | 0.38 |
| 75 days, organic | 75 days, conventional | 1.21 | 0.21 | 1.09 | 0.39 |
| 75 days, organic | 45 days, conventional | 1.45 | 0.26 | 2.08 | 0.07 |
| 75 days, organic | 105 days, conventional | 1.02 | 0.18 | 0.11 | 0.91 |
| 45 days, conventional | pre-b, conventional | 2.64 | 0.40 | 6.40 | **<0.0001** |
| 45 days, conventional | post-b, conventional | 1.10 | 0.13 | 0.77 | 0.55 |
| 45 days, conventional | 75 days, conventional | 0.84 | 0.09 | –1.60 | 0.18 |
| 45 days, organic | pre-b, conventional | 3.97 | 0.81 | 6.80 | **<0.0001** |
| 45 days, organic | pre-b, organic | 1.40 | 0.18 | 2.63 | **0.02** |
| 45 days, organic | post-b, conventional | 1.65 | 0.30 | 2.79 | **0.01** |
| 45 days, organic | post-b, organic | 0.91 | 0.11 | –0.80 | 0.55 |
| 45 days, organic | 75 days, conventional | 1.26 | 0.22 | 1.32 | 0.28 |
| 45 days, organic | 75 days, organic | 1.04 | 0.12 | 0.33 | 0.78 |
| 45 days, organic | 45 days, conventional | 1.50 | 0.27 | 2.30 | **0.04** |
| 45 days, organic | 105 days, conventional | 1.06 | 0.18 | 0.33 | 0.78 |
| 105 days, conventional | pre-b, conventional | 3.75 | 0.55 | 9.03 | **<0.0001** |
| 105 days, conventional | post-b, conventional | 1.56 | 0.17 | 3.99 | **<0.0001** |
| 105 days, conventional | 75 days, conventional | 1.19 | 0.12 | 1.69 | 0.16 |
| 105 days, conventional | 45 days, conventional | 1.42 | 0.15 | 3.26 | **<0.0001** |
| 105 days, organic | pre-b, conventional | 4.11 | 0.83 | 6.99 | **<0.0001** |
| 105 days, organic | pre-b, organic | 1.45 | 0.19 | 2.89 | **0.01** |
| 105 days, organic | post-b, conventional | 1.70 | 0.30 | 2.98 | **0.01** |
| 105 days, organic | post-b, organic | 0.94 | 0.11 | –0.51 | 0.69 |
| 105 days, organic | 75 days, conventional | 1.30 | 0.23 | 1.52 | 0.20 |
| 105 days, organic | 75 days, organic | 1.08 | 0.13 | 0.62 | 0.65 |
| 105 days, organic | 45 days, conventional | 1.56 | 0.27 | 2.50 | **0.03** |
| 105 days, organic | 45 days, organic | 1.03 | 0.12 | 0.29 | 0.79 |
| 105 days, organic | 105 days, conventional | 1.10 | 0.19 | 0.53 | 0.69 |
